# Immune disease variants modulate gene expression in regulatory CD4+ T cells and inform drug targets

**DOI:** 10.1101/654632

**Authors:** Lara Bossini-Castillo, Dafni A. Glinos, Natalia Kunowska, Gosia Golda, Abigail Lamikanra, Michaela Spitzer, Blagoje Soskic, Eddie Cano-Gamez, Deborah J. Smyth, Claire Cattermole, Kaur Alasoo, Alice Mann, Kousik Kundu, Nicole Soranzo, Ian Dunham, David Roberts, Gosia Trynka

## Abstract

Identifying cellular functions dysregulated by disease associated variants could implicate novel pathways for drug targeting or modulation in cell therapies. Variants associated with immune diseases point towards the role of CD4+ regulatory T cells (Tregs), cell type critical for immune homeostasis. To translate the effects of immune disease alleles on Treg function we mapped genetic regulation (QTL) of gene expression and chromatin activity in Tregs. We identified 123 loci where Treg QTLs colocalized with immune disease variants. These colocalizations indicated that dysregulation of key Treg pathways, including cell activation via CD28 and IL-2 signalling mediated by STAT5A contribute to the shared pathobiology of immune diseases. Finally, using disease GWAS signals colocalizing with Treg QTLs, we identified 34 known drug targets and 270 targets with drug tractability evidence. Our study is the first in-depth characterization of immune disease variant effects on Treg gene expression modulation and dysregulation of Treg function.

## Introduction

Thousands of disease variants mapped through genome wide association studies (GWAS) provide genetic signposts to disease pathophysiological pathways but functional interpretation of GWAS signals has been challenging as the vast majority of variants are non-coding. One approach for linking genetic variation to regulated genes includes expression quantitative trait locus (eQTL) mapping, in which transcript levels are correlated with genetic polymorphisms^1^. However, due to the linkage disequilibrium (LD) between genetic variants, the identified eQTLs often result in associations of tens to hundreds of correlated variants with gene expression levels, and therefore fail to nominate the causal regulatory variants.

Prioritization of the exact regulatory variants underlying gene expression changes can be further inferred through QTL mapping of chromatin activity using chromatin accessibility or histone modifications (chromatin QTLs; chromQTLs). In this approach, variants that are linked to modulate the levels of chromatin marks can be physically overlapped with the chromQTL features^2^. The combination of eQTLs and chromQTLs is a powerful tool for linking non-coding variants to genes whose expression is modulated, for prioritizing functional variants and for identifying mechanisms through which gene expression is regulated. Finally, colocalization^3^ of disease GWAS signals with such QTLs can point towards causal genes and mechanisms underlying disease associations, therefore linking disease associated variants to dysregulated pathways and new drug targets.

GWAS variants associated with common immune-mediated diseases such as inflammatory bowel disease (IBD), type 1 diabetes (T1D) and rheumatoid arthritis (RA) are enriched in active chromatin marks that tag enhancers and promoters in the CD4+ T cell compartment, especially in regulatory T cells (Tregs)^4–6^. Tregs are an infrequent yet functionally significant subset of CD4+ T cells, they comprise 2-10% of CD4+ T cells and play an essential homeostatic role in the immune system by suppressing the proliferation and effector functions of conventional T cells. Immunophenotyping studies have shown that abnormal numbers of circulating Tregs^7,8^ and defective suppressive function of Tregs results in a dysregulated immune response in patients with immune diseases^9–11^, as well as in organ and hematopoietic stem cell transplant recipients ^12,13^. Therefore, identifying how variants modulate Treg function could have important clinical implications. Already, *ex vivo* approaches to expand Treg numbers and to enhance Treg suppressive capacity and reinforce them into patients with immune diseases have been successful in clinical trials for type 1 diabetes^14–16^ and Crohn’s disease^17^.

Despite the key role of Tregs in maintaining appropriate immune responses and due to their low frequency, there are only limited large scale genomic resources available ^18–20^. Consequently, immune disease variants are often interpreted in light of gene expression data from peripheral blood mononuclear cells (PBMCs), immune cell lines or isolated major immune cell populations ^2,21–26^. However, these datasets can either dilute or omit gene regulatory effects only present in scarce cell types, therefore potentially missing biological effects meaningful to the disease.

Here, we generated a detailed resource of gene expression regulation in Tregs to interpret immune disease variants in light of Treg biology. We identified a total of 14,313 QTL effects (4,133 eQTLs and 10,180 chromQTLs), with 32% of the eQTLs and 57% of the active enhancer and promoter QTLs specific to Tregs and undetected in the closely related naive CD4+ T cells ^24^. By colocalizing Treg eQTLs with 14 immune disease associated variants we assigned candidate functional genes to 61 immune disease loci. The overlap of immune disease GWAS signals with chromQTLs functionally refined associated variants at 66 immune disease loci. Finally, we used the prioritised genes to identify drugs for repurposing and to define novel targets for validation. Our study provides a translational pathway from immune disease associated variants, through gene expression regulation in Tregs, to new treatment options.

## Results

### Comprehensive catalogue of gene expression regulation in Tregs

To identify genetic variants that control gene expression regulation in Tregs isolated from healthy blood donors (**Supplementary Figure 1**) we profiled the transcriptome using RNA-seq (123 individuals), chromatin accessibility using ATAC-seq (73 individuals), promoters using H3K4me3 (88 individuals), and active enhancer and promoter regions using H3K27ac (91 individuals; Figure 1A). We detected the expression of 12,059 genes, while chromatin profiling revealed 39,642 accessible regions, 40,285 H3K4me3 marked promoter regions, and 34,457 H3K27ac marked active chromatin regions (Figure 1B). The majority of the mapped regulatory chromatin features overlapped with each other (**Supplementary Figure 2A**). Concordant with previous studies ^24,27,28^, we observed that H3K4me3 and chromatin accessible regions were concentrated near the transcription start sites (TSS), while H3K27ac marked more distal gene regulatory elements (**Supplementary Figure 2B**).

**Figure 1.**
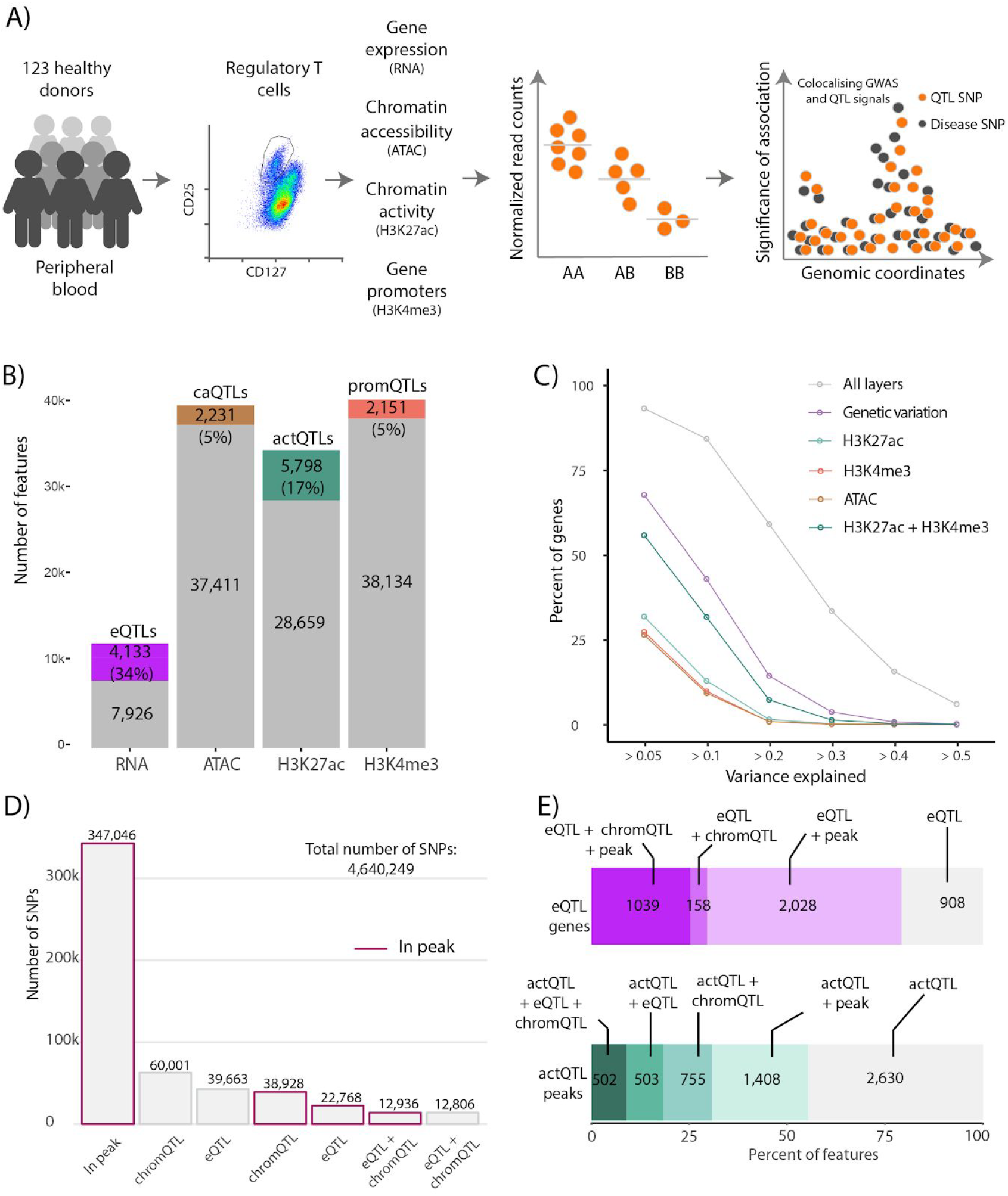
Overview of mapped Treg QTLs. **A)** A schematic of our study design. **B)** Number of features defined per functional layer and number of significant QTLs in each category. **C)** Proportion of gene expression variance explained by genetic variation and chromatin marks. We considered cis-regulatory elements in a ± 150kb window from the gene. Shown is the cumulative contribution of genes with increasing proportions of explained variance. **D)** Functional classification of tested genetic variants. Bars with purple outline indicate instances when a QTL variant maps to any chromatin peak. Categories are mutually exclusive. **E)** Classification of eQTL genes (top) and actQTL peaks (bottom). eQTL genes were classified based on the annotation of eQTL variants with chromQTLs and overlap with chromatin peaks. eQTL+chromQTL+peak, number of eQTL genes for which eQTL variants also result in a chromQTL and one of the eQTL variants mapped within a chromatin mark peak; eQTL+chromQTL, number of eQTL genes for which eQTL variants also result in a chromQTL but no variant mapped within any chromatin mark peak; eQTL+peak, number of eQTL genes for which eQTL variants map within a chromatin mark peak but no chromQTL effects were detected; eQTL, number of eQTL genes for which we were unable to map eQTL variants to chromatin mark peaks or to link them to chromQTLs. actQTL peaks were classified based on the annotation of actQTL variants with eQTLs and overlap with chromatin peaks. actQTL + eQTL + chromQTL, number of actQTLs that also result in an eQTL and an additional chromQTL; actQTL + eQTL, number of actQTLs that also result in an eQTL; actQTL + chromQTL, number of actQTLs which also result in an additional chromQTL; actQTL + peak, number of actQTLs that map within a chromatin mark peak without an additional chromQTL or eQTL; actQTL, number of actQTLs that we were unable to map variants to chromatin mark peaks or to link them to an additional chromQTL or eQTL.

Using the 62 samples for which we had complete information including genetic variation, chromatin profiles and whole transcriptome, we estimated the percentage of gene expression variability explained by the genetic component and by the chromatin regulatory features. We observed that the major component driving transcriptional variability was common genetic variation, which explained 5% or more of the gene expression variance of nearly 68% of genes (Figure 1C). The combination of chromatin marks (H3K27ac, H3K4me3 and ATAC) with common genetic variation explained 5% or more of the gene expression variance for an additional 17% of genes. This additional gene expression variability was mainly accounted for by the combination of H3K27ac and H3K4me3. Together, the gene expression variance decomposition analysis implicated that genetic variation contributed the most towards gene expression regulation, and the genetic effects manifested both on the transcriptome and the chromatin regulatory marks. Therefore, by connecting genetic variation to gene expression and chromatin regulatory features we expected our Treg dataset to provide translational insight into immune disease GWAS loci.

Next, we performed QTL mapping to define genes and chromatin features that were under genetic control in Tregs (FDR < 0.05; see **Methods**; ^29^). We detected at least one independent association for 4,133 genes (34%) with a total of 88,173 variants (eQTLs; Figure 1B). We mapped a total of 10,180 chromQTLs, using chromatin accessibility (caQTLs, 2,231; 5%), H3K4me3 (promQTLs, 2,151; 5%) and H3K27ac (actQTLs, 5,798; 17%) histone marks, which corresponded to 7,652 unique regions, associated with 124,671 variants. The majority of chromQTLs were actQTLs, where we detected 5,798 regions under genetic control (Figure 1B). Of all analyzed genetic variants (4,640,249) only a small fraction mapped in at least one chromatin feature and was either linked to chromQTLs (38,928; 0.84%), eQTLs (22,768; 0.49%) or both (12,936; 0.28%) (Figure 1D). For 25% (1,039) of all eQTL genes we observed that at least one eQTL variant was a chromQTL and was also physically located in a chromatin peak (Figure 1E), and for an additional 2,028 eQTL genes we were able to link an eQTL variant to a chromatin peak however we did not detect a QTL effect on any chromatin feature. A proportion of this overlap may not be functional as chromatin regulatory features are abundant throughout the genome and therefore likely to overlap common genetic variants by chance. Interestingly, we were unable to link the majority of actQTL variants to an eQTL (Figure 1E), implicating that these regulatory regions may modulate gene expression under a specific cellular context, or through the interplay of multiple regulatory elements.

### Treg QTLs colocalize with immune-disease loci and functionally refine GWAS associations

We first sought to estimate the proportion of QTLs specific to Tregs and absent from closely related naive CD4+ T cells. For this, we retrieved both RNA and H3K27ac profiles from purified CD4+ naive T cells ^24^ and compared the eQTL and actQTL signals between the two cell types. As expected, we observed that the majority of eQTLs were shared (68%; Figure 2A) with similar effect sizes and the same direction of effects (**Supplementary Figure 5A**; Spearman correlation = 0.86). However, examination of the eQTLs and actQTL signals in Tregs, showed evidence of Treg-specific modulation of gene expression. We classified 1,309 genes (32% of all eQTL genes) as Treg specific eQTLs, including 167 genes that were only expressed in Tregs (see **Methods)**. Among the Treg specific eQTLs, there were many genes related to immune function, including *CCL20* (FDR=3.87×10^−3^), a chemokine that attracts lymphocytes towards epithelial cells, and *CD28* (FDR=4.37×10^−11^), a co-stimulatory receptor critical for activation of T cells ^30^ (Figure 2B and **Supplementary Figure 3**). However, when we compared the actQTLs called in both cell types we observed that 3,326 (57%) actQTLs were Treg specific. For example, the same variant (chr2:227,807,863; rs13034664) that affected the expression of *CCL20* also regulated the activity of a nearby enhancer (chr2:227,804,673-227819641; Figure 2C). This enhancer peak was absent in naive T cells. We observed that of the 167 eQTL genes that were only expressed in Tregs, 53 had a significant Treg actQTL that was controlled by the same variant (or a variant in strong LD).

**Figure 2.**
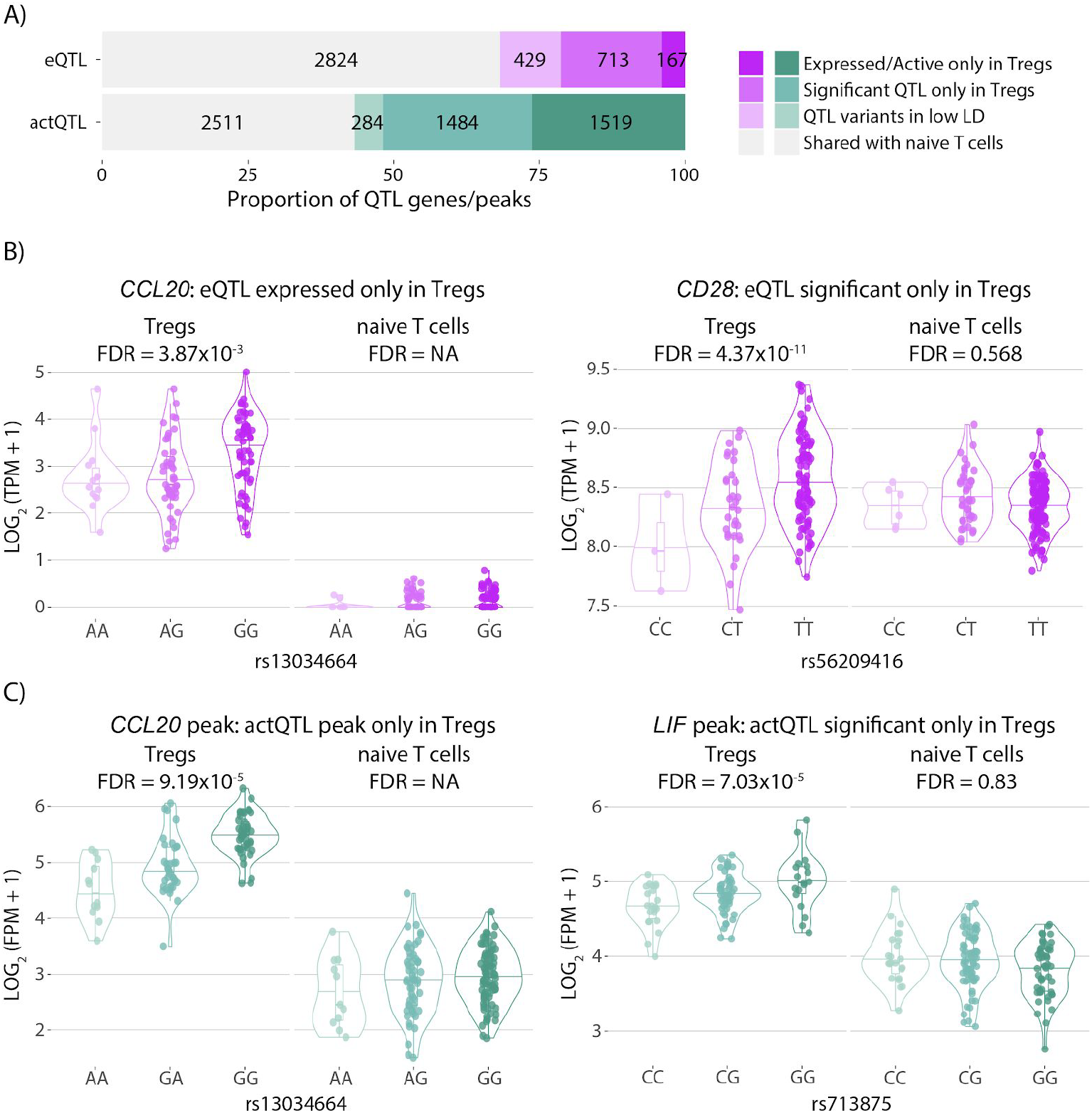
Comparison of eQTLs and actQTLs identified in CD4+ naive and regulatory T cells. **A**) Proportion of eQTLs and actQTLs specific to Tregs in comparison to naive T cells. **B**) Examples of Treg specific eQTLs and **C)** actQTLs.

To fine-map disease associated loci to causal genes and variants we next integrated the Treg QTL results with GWAS signals from common immune diseases. We applied a Bayesian framework to test for statistical colocalization of the disease associated variants and the Treg QTL signals ^31^. Collectively, we tested 1,290 unique GWAS loci associated with 14 immune mediated diseases: allergic diseases (ALL), ankylosing spondylitis (AS), asthma (AST), celiac disease (CEL), Crohn’s disease (CD), inflammatory bowel disease (IBD), multiple sclerosis (MS), primary biliary cirrhosis (PBC), psoriasis (PS), rheumatoid arthritis (RA), systemic lupus erythematosus (SLE), type 1 diabetes (T1D), ulcerative colitis (UC) and vitiligo (VIT) (see **Methods**; Figure 3A). Diseases with the highest number of colocalizations (more than 20 colocalizing signals) included IBD, UC, CD, ALL and RA. The high number of observed colocalizations is consistent with previous work that implicated the role of Tregs in the pathobiology of all of these diseases ^5,6,32–34^, as well as the higher number of significant GWAS loci for these traits. Overall, immune-mediated diseases showed more colocalizations with Treg QTLs than non-immune mediated diseases, such as type-2 diabetes or depression (Figure 3A).

**Figure 3.**
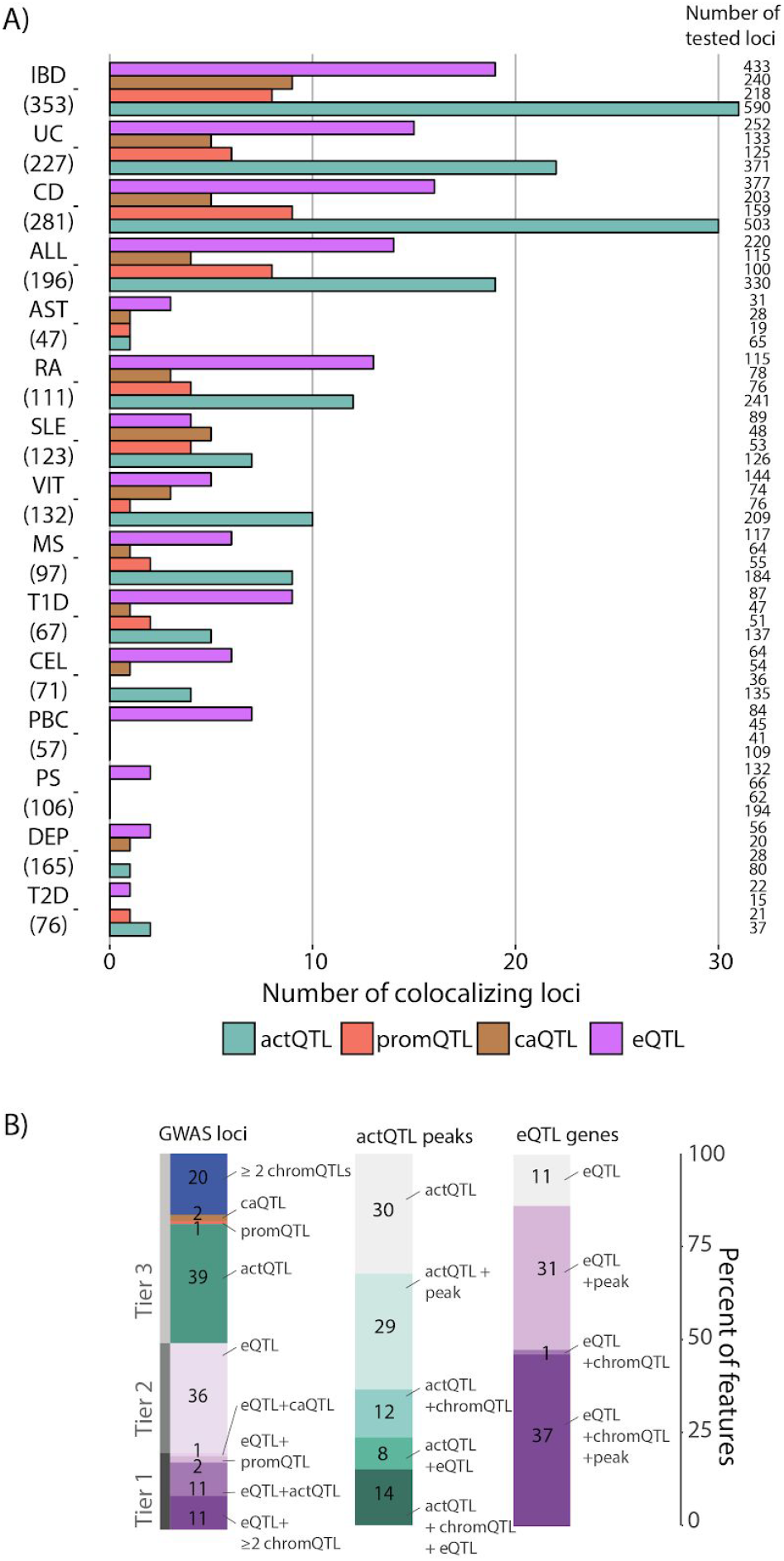
Colocalization of immune disease GWAS loci and Treg QTLs. **A)** Distribution of Treg eQTLs and chromatin QTLs colocalizing with different immune disease GWAS loci. Number in parenthesis is state independent loci associated with the trait. The numbers on the right side of the bars correspond to the total number of features (genes or peaks) tested for colocalization. ALL: allergic disease (asthma, hay fever and eczema); AST: asthma; CD: Crohn’s disease; CEL: celiac disease; DEP: Broad depression; IBD: inflammatory bowel disease; MS: multiple sclerosis; PBC: primary biliary cirrhosis; PS: psoriasis; RA: rheumatoid arthritis; SLE: systemic lupus erythematosus; T1D: type-1 diabetes; T2D: type-2 diabetes; UC: ulcerative colitis; VIT: vitiligo. **B)** Distribution of the GWAS loci colocalizing with different types of Treg QTLs.

We observed 240 significant colocalizations between the disease loci and at least one Treg QTL (Figure 3A, **Supplementary Table 1-2**), corresponding to 123 unique GWAS loci. Of the 123 unique GWAS loci, 36 loci colocalized with eQTLs, 62 with chromQTLs, and 25 colocalized with both eQTL and at least one chromQTL (Figure 3B). These colocalizations affected the expression of 80 eQTL genes (Figure 3B), acetylation of 93 actQTL peaks (Figure 3B), methylation of 31 promQTL peaks and accessibility of 25 caQTL sites. Four of the colocalizing eQTLs were shared between four or more diseases and included *BACH2* (ALL, AST, CEL, IBD, T1D, and VIT), *GSDMB* and *ORMDL3* (ALL, AST, IBD, PBC, RA, SLE and T1D) and *TYK2* (PBC, PS, RA, SLE and T1D). While we observed the largest number of colocalizations with Treg actQTLs, there was only one actQTL, (chr11:76,586,431-76,600,121) near *LRRC32*, that colocalized with four diseases (ALL, AST, IBD and T1D) (**Supplementary Table 1**). Of the immune disease GWAS loci which colocalized with both Treg eQTLs and chromQTLs, 21 out of 25 were actQTLs (84%) (Figure 3B). Finally, for the vast majority (83.5%) of the loci where we observed disease signals colocalizing with two or more types of QTLs, the effects of the risk alleles propagated in the same direction across the different functional layers. For example, the *IL7* eQTL colocalized with MS variants chr8:78663569 and the risk alleles resulted in both reduced gene expression and decreased H3K27 acetylation, H3K4 tri-methylation and chromatin accessibility (**Supplementary Table 1**). However, at 14 loci we observed that the disease alleles resulted in opposite effects between the different types of QTLs. For example, the MS risk allele, chr11:71,457,027-A (rs4944958), increased the expression of *DHCR7* as well as the acetylation of the chr11:71,446,806-71,463,563 actQTL peak while it decreased the expression of *NADSYN1* (**Supplementary Table 1**; **Supplementary Figure 4**). The MS risk association signal colocalized with chr11:71,449,349-G (rs4944949), the lead variant for the three QTLs described above, and resulted in increased binding of FOXP1, a transcriptional repressor in immune cells ^35,36^. These cases suggest complex mechanisms of gene expression regulation.

We systematically investigated all immune disease signals colocalizing with Treg QTLs to refine the disease associated signals to sets of functional variants and to nominate causal genes. We classified colocalizing loci into three categories. Tier 1 loci comprised 25 signals for which the GWAS association colocalized with both eQTL and chromatin QTL (Figure 3B). Of these, at 20 loci the associated variants were also located within the chromQTL peaks. Loci in this category were the most informative to functionally refine disease associations, as we were able to link the GWAS signals to genes and to functional chromatin elements that regulated gene expression. Tier 2 loci contained 36 signals for which we observed colocalization only with eQTLs (Figure 3B). In this case we were unable to refine the association signals to sets of functional variants but we were able to connect the GWAS signals to candidate causal genes. Finally, the 62 loci in Tier 3 included GWAS signals colocalizing with chromatin QTLs but not eQTLs. Of these, at 44 loci the GWAS variants overlapped a chromatin QTL peak, providing further clues to prioritize functional variants at GWAS loci. Finally, Tier 3 loci represented the majority of colocalizations. We hypothesized that gene expression effects could be manifested in a cell state specific context. To further nominate candidate genes regulated by the variants colocalizing with actQTLs, we used resting and activated Treg transcriptome data (see Methods) and defined genes proximal to the QTL peaks that were differentially expressed upon cell activation (**Supplementary Table 1**). This analysis prioritized 72 genes linked to 32 disease colocalizing actQTLs. Furthermore, we used CAGE data from FANTOM5 (Andersson et al., 2014) and physically linked 40 of the disease colocalizing actQTLs to 231 genes (see Methods). Of these, 22 actQTLs were connected to 32 genes differentially expressed upon Treg stimulation including *AIRE*, *CD247*, *CXCR5*, *LRRC32* and *PRDM1*.

Together, using the Tier 1 and Tier 3 loci that overlapped with chromatin QTL peaks, we refined the signals at 66 GWAS loci from an average of 79 associated variants to 9 functional variants per locus (Figure 4A). Of the 66 loci, in 38 instances we observed that the genetic variants additionally overlapped open chromatin peaks allowing us to further prioritize the functional variants from an average of 9 functional variants to an average of 4 variants per locus, including *BACH2*, *CD28*, *GBA*, *GSDMB*/*ORMDL3*, *IL7*, *MAP3K8*, *PIM3*, *STAT5A* and *TNFRSF9* loci which colocalized with eQTLs and we refined to a single functional variant (Figure 4B and **Supplementary Table 3**). In the case of previously statistically fine-mapped loci, in which associations have been refined to rare variants or haplotypes, such as *CD28*, *BACH2*, *CTSH* and *TYK2* ^37,38^, the information from Treg QTL colocalizations prioritized additional functional variants.

**Figure 4.**
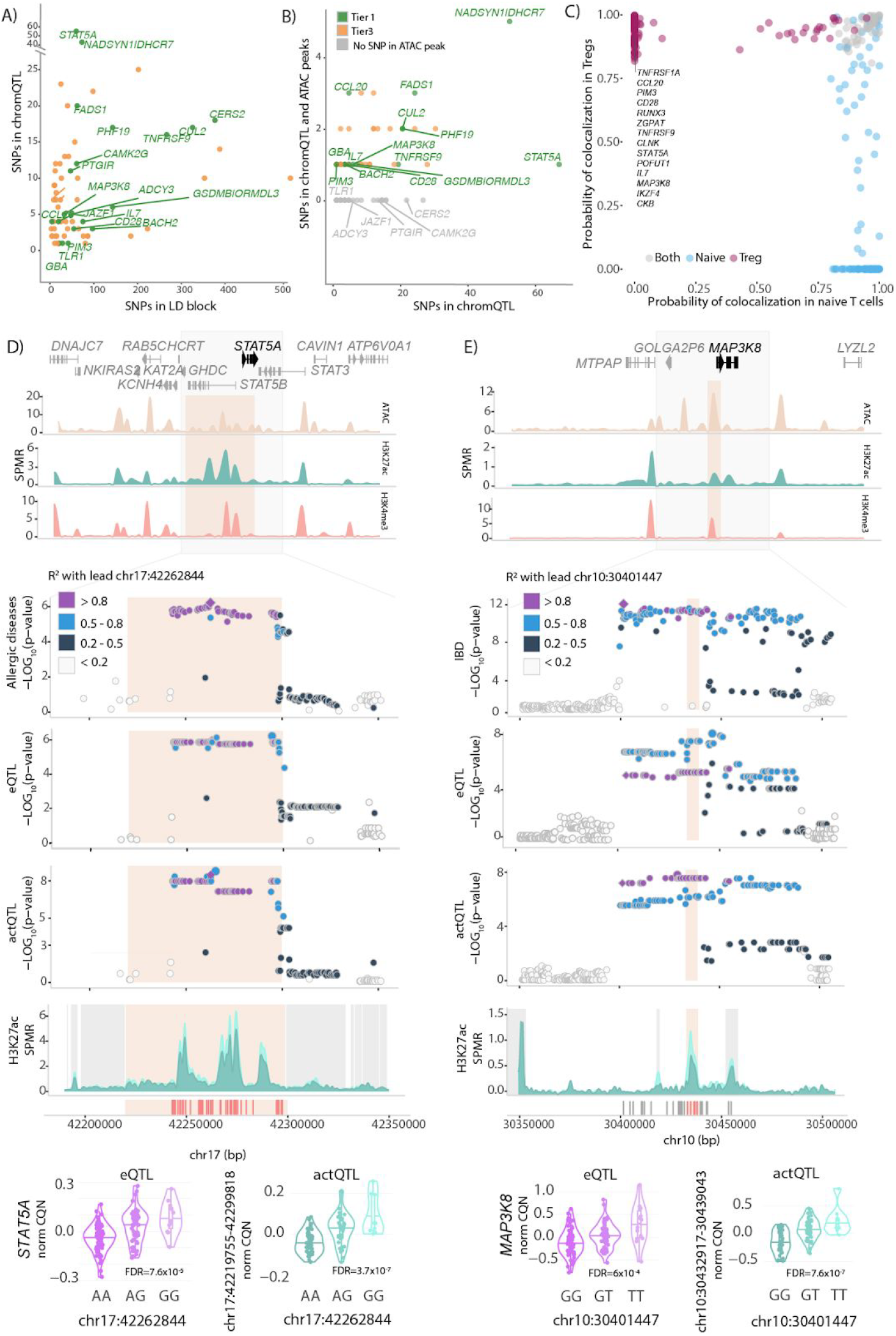
Functional refinement of immune disease associations colocalizing with Treg QTLs. **A)** The number of SNPs in LD blocks (lead GWAS signals and their proxies R^2^ ≥ 0.8) on the x-axis, and the number of SNPs that map inside chromQTL peaks on the y-axis. **B)** The number of SNPs in LD blocks that map inside chromQTL peaks on the x-axis and the number of SNPs that map inside both chromQTL and an additional ATAC peak on the Y axis. **C)** Shared and cell type specific *coloc* values of immune disease GWAS signals colocalizing with eQTLs mapped in naive and regulatory T cells. *Coloc* values were calculated as PP.H4/(PP.H4+PP.H3). Genes are colored based on colocalization in either or both cell types. Labels indicate Treg specific eQTLs. **D)** and **E)** panels. From top to bottom the figure displays: gene annotation tracks, chromatin landscape for ATAC-seq, H3K27ac and H3K4me3 ChM-seqs, region association plots for disease, eQTL and actQTL association p-values, focussed on H3K27ac landscape stratified by homozygous genotypes and genotype stratified eQTL and actQTL violin plots. **D)** Locus associated with allergies, tagged by chr17:42,262,844 (rs7207591) SNP colocalizing with *STAT5A* eQTL and chr17:42,219,755-42,299,818 actQTL. **E)** Locus associated with IBD, tagged by chr10:30,401,447 (rs10826797) SNP colocalizing with *MAP3K8* eQTL and chr10:30,432,917-30,439,043 actQTL.

Next, we assessed which of the identified eQTLs that colocalized with immune disease variants regulated gene expression specifically in Tregs and not in naive T cells. Out of the 61 GWAS loci that showed colocalization with eQTLs 22 were Treg exclusive, i.e. were not present in naive T cells (Figure 4C). To ensure that this was not due to technical issues, e.g. poorly captured transcripts in naive T cells, we further removed the genes for which we observed eQTL effects in five or more GTEx tissues ^26^ (see **Methods**). This resulted in a final set of 14 Treg exclusive eQTLs that colocalized with immune disease GWAS variants (Figure 4C and **Supplementary Figure 5B**): *CCL20*, *CD28*, *CKB*, *CLNK*, *IKZF4*, *IL7*, *MAP3K8*, *PIM3*, *POFUT1*, *RUNX3*, *STAT5A*, *TNFRSF1A*, *TNFRSF9* and *ZGPAT*. For eight of these eQTLs we also found a colocalization with a Treg chromQTL: *CCL20*, *CD28*, *CLNK*, *IL7*, *MAP3K8*, *PIM3*, *STAT5A*, and *TNFRSF9*.

The Treg exclusive colocalizations indicated regulation of pathways that were characteristic of Treg biology. For example, we observed that a locus associated with allergies ^39^ (tagged by the index SNP chr17:42,262,844, rs7207591) colocalized with a *STAT5A* eQTL (p-value = 5.93 × 10^−7^), as well as with an 80 kb actQTL (chr17:42,219,755-42,299,818) (Figure 4D). This peak overlapped the *STAT5A* transcription start site (TSS). Nearly the entire LD block of allergy variants (55 SNPs) overlapped with the regulated actQTL peak and one of the variants, chr17:42,266,938 (rs34129849), also mapped to a 629 bp open chromatin region (chr17:42,266,595-42,267,224) located in intron 1 of the *STAT5B* gene (Figure 4D, **Supplementary Table 1** and **3**). Modulation of STAT5 mediated pathways could implicate broad effects on Treg function as STAT5A regulates the expression of genes downstream of the IL-2 receptor which is critical for Treg development and function ^40^.

Another example of a Treg exclusive colocalization included an IBD GWAS signal, tagged by the chr10:30,401,447 (rs10826797) variant, which colocalized with an actQTL, regulating a 6kb large (chr10:30,432,917-30,439,043) H3K27ac peak (p-value = 1.97 × 10^−9^) at the TSS of *MAP3K8*, and an eQTL (p-value = 6.79 × 10^−9^) for the *MAP3K8* gene (Figure 4E and **Supplementary Table 1**). The IBD risk allele increased the acetylation at H3K27 and upregulated the expression of *MAP3K8*. Five of the colocalizing variants overlapped this actQTL peak, of which only one SNP, chr10:30,434,664 (rs306588), overlapped a 1.5 kb ATAC peak (chr10:30,433,210-30,434,733) (**Supplementary Table 1** and **3)**. This approach refined the IBD associated signal from 30 GWAS variants to a single functional candidate variant regulating the expression of *MAP3K8*, a kinase modulating the DNA binding activity of FoxP3, the Treg hallmark transcription factor ^41^.

### Treg QTLs emphasize CD28 co-stimulation, TNF and IL-10 signaling pathways for drug targeting

Despite the success of GWAS in mapping disease risk variants the efforts to translate these findings into drug targets have been challenging. Therefore, we systematically assessed if eQTLs that colocalized with immune disease signals could identify potential drug targets (see **Methods**). We found that ten Treg eQTLs (Tier 1: *CD28, GBA, PIM3, PTGIR, TNFRSF9* and Tier 2: *BLK, ERAP2, NDUFS1, TNFRSF1A, TYK2*) (Figure 5A) and 5 genes prioritized in Tier 3 loci (*CDK4, PDCD1, PRKCD, PSMA6, and TNNT3*) (**Supplementary Table 1, 2** and **4**) were already targeted by known drugs and were either used in clinical practice or undergoing clinical trials. Most of these targets are not exclusively expressed in Tregs, but our characterization of the cell type specific effects of increased or decreased gene expression on disease risk suggests that targeting these pathways alone, or in combination with other Treg modifying agents might have therapeutic benefits.

**Figure 5.**
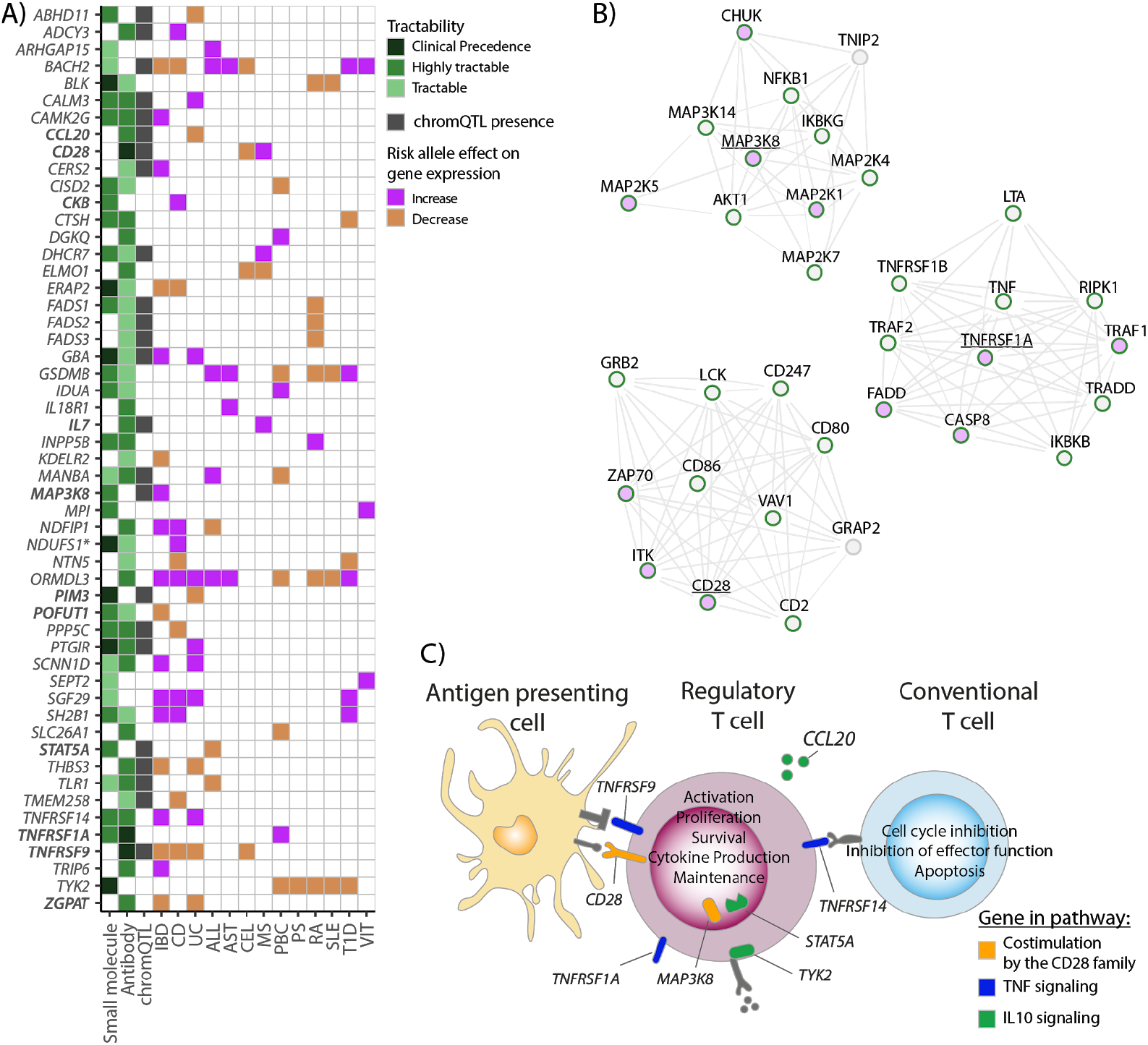
Immune disease colocalizations with Treg QTLs inform drug targets. **A)** Tier 1 and Tier 2 loci colocalizing with immune disease GWAS variants with drug tractability evidence (green). In bold are Treg specific eQTLs. Clinical precedence: gene targeted by small molecules or antibodies approved for patient treatment or undergoing clinical trials; Discovery precedence: gene product shown to bind small molecules; Tractable high confidence: gene product with high predicted tractability as an antibody drug target; Tractable medium - low confidence: gene product with predicted tractability as an antibody drug target. Predicted tractable: gene predicted to be small molecule-tractable. *For *NDUFS1*, the encoded protein is not directly targeted but is part of a targeted complex. **B)** Examples of physical and functional protein-protein interactions for immune disease colocalizing Treg eQTLs (underlined). Genes with tractability evidences are depicted with green borders, Treg eQTLs are in purple. **C)** Tier 1 and Tier 2 genes with tractability potential in CD28-costimulation (orange), TNF (blue), and anti-inflammatory IL-10 (green) pathways.

We identified five eQTL genes that colocalized with immune disease GWAS variants which could be considered for drug repurposing: *NDUFS1*, *TNFRSF1A*, *TNFRSF9*, *TNFRSF14* and *TYK2*. For example, a colocalization between a *NDUFS1* eQTL and CD, in which the disease risk allele increased gene expression, suggested repurposing metformin which targets the NADH dehydrogenase complex (not directly *NDUSF1*). Metformin is currently used for treating type 2 diabetes and in a cohort of MS patients increased the number of Tregs ^42^. On the other hand, we observed eQTLs that colocalized across multiple diseases but with opposite allelic effects, therefore arguing against drug repurposing. For example, both CEL and MS colocalized with *CD28* eQTL but the CEL risk allele led to decreased *CD28* expression, while the MS risk allele increased gene expression (Figure 5A). Finally, we observed 72 genes that were not yet a part of a clinical treatment but had drug tractability evidence, of which 56 were classified as highly tractable (Figure 5A, **Supplementary Table 1, 2** and **4**). One such example was *MAP3K8*, for which there are three ChEMBL compounds that target it. Importantly, we observed that the IBD risk allele colocalizing with the *MAP3K8* eQTL increased gene expression, implicating *MAP3K8* as a good target for validation.

Networks that include genes for which disease risk alleles increase gene expression could be particularly informative for target validation, for example, we observed that physical and functional protein-protein interaction networks built for *MAP3K8*, *TNFRSF1A* and *CD28* included multiple genes with tractability evidence (Figure 5B). Collectively, we observed that genes with high tractability evidence fell into co-stimulation by the CD28 family, TNF signaling, and IL10 signaling pathways (Figure 5C). These pathways play an important role in Treg cell activation, proliferation and survival, as well as in suppression of effector T cells.

## Discussion

Pinpointing genes that are regulated by disease associated non-coding variants can uncover important cell pathways for drug targeting. However, leveraging information captured by GWAS variants to provide insight into disease biology and improve treatment has been challenging. Increasing availability of functional genomic resources from different cell types helps to bridge this gap. Naive and regulatory CD4+ T cells are closely related, yet they play distinct functions in the immune system. Although naive CD4+ T cells have been extensively characterized, Tregs are an infrequent cell population difficult to isolate in large numbers for QTL analysis and elusive to deconvolute from bulk blood QTL data ^43^. Therefore, mapping gene expression regulation directly in Tregs is essential to better understand Treg biology.

Here, we sought to describe the role of immune disease associated variants on modulation of gene expression in Tregs. We linked 123 unique immune disease loci from associated variants to functional effects; 36 loci were linked to gene expression, 62 loci were linked to an effect on chromatin and 25 loci to both. Loci for which we observed colocalization with both gene expression and chromatin QTLs provide an important translational insight into mechanisms through which immune disease variants regulate Treg function. For example, we observed signals overlapping with Treg specific eQTLs, indicating regulation of essential Treg pathways, such as IL-2 signalling via *STAT5A*.

The 62 loci for which we only detected colocalization with chromQTLs but not eQTLs indicate that the altered gene expression may be manifested in a specific cell state, which will require tailored functional follow up studies. Indeed, previous studies showed that context specific eQTLs, e.g. stimulation induced, can be already detected in a resting state at the chromQTL level ^2,22,24^. A subset of loci where we were unable to link chromQTLs with eQTLs could also result from long distance gene expression regulation, the eQTL being outside of our testing window, or from combinatorial subtle effects between multiple enhancers regulating the expression of individual genes. We also recognize instances of complex regulation of gene expression that will require targeted follow-up studies to fully uncover the functional role of disease variants. For example, we observed complex pattern of colocalization between *CD28* eQTL, nearby actQTLs and immune disease GWAS variants (**Supplementary Figure 3**). The eQTL for *CD28*, the co-stimulatory receptor found on the surface of the majority of T cells, was specific to Tregs and absent from naive T cells. The risk alleles for CEL and MS showed reversed effects on *CD28* expression and the acetylation of the peaks, implicating complex enhancer-mediated control of *CD28* expression under cell type and cell state specific mechanisms.

On the other hand, the 36 loci colocalizing only with eQTL variants but not chromQTLs may be correlated with chromatin-independent gene expression regulation, such as splicing QTLs (sQTLs)^44^ or RNA stability^45^. For example, we observed that *ERAP2*, an IBD and CD associated locus, showed an eQTL colocalization but no chromQTL effect. The lead GWAS variant chr5:96,912,106 (rs6873866) and the colocalizing lead eQTL variant chr5:96,916,728 (rs2927608) are proxies for chr5:96,900,192 (rs2248374), a sQTL present in monocyte-derived dendritic cells after influenza infection and type 1 interferon stimulation^46^.

By linking immune disease GWAS variants to Treg eQTLs our study provides genetic evidence for discovery and repurposing of drugs that modulate Treg function in treating immune disease patients. Validation of targets with genetic support can significantly increase the chance of clinical success^47,48^. Particularly, our results support the focus on modulating costimulatory and cytokine pathways, and highlight the importance of developing cell type specific drug targeting approaches. For example, we observed a shared association between IBD, UC and CEL, in which the disease risk alleles led to decreased levels of expression of *TNFRSF9* (encodes for CD137/4-1BB). Signaling via CD137 induces cell division and proliferation^49,50^, however, *TNFRSF9* gene expression and protein levels increase specifically in activated Tregs but not in conventional T cells^51,52^. Furthermore, the increased expression of CD137 enhances the Treg capacity to suppress proliferation of effector T cells^53^. Therefore, the disease risk allele could result in decreased Treg suppressive function and promote immune disbalance. Agonistic anti-TNFRSF9 monoclonal antibody therapy, which enhances the CD137 signalling pathway, has had effective results in the treatment of patients with inflammatory bowel disease^54^. The colocalization of several TNF receptor superfamily members (*TNFRSF1A*, *TNFRSF9*, *TNFRSF14*) further supports the development of improved anti-TNF biologics, one of the main therapy lines for treating immune diseases, and extends the list of diseases they can be used for (Figure 5A).

Understanding the genetic underpinnings of immune system regulation will have broad implications not only in the treatment of immune mediated conditions, but also in infections, transplantation, and cancers. For instance, in organ transplantation, numbers of Tregs, as well as Tregs with increased suppressive capacity can provide a favourable environment of successful transplant tolerance^12,55,56^. Furthermore, in haematopoietic stem cells transplantation high Treg:CD4 T cell ratios are associated with reduced acute graft-versus-host disease and reduced overall mortality^56^. Importantly, *in vitro* expanded Tregs with enhanced suppressive capacity have already entered clinical trials^57^. Therefore, identifying genetic variants that regulate gene expression in a specific cellular context can inform development of more effective cell therapies. Our study provides an important advancement in mapping regulation of gene expression in Tregs and consequently our results can benefit a range of clinical conditions.

## Acknowledgements

We thank all participating blood donors and the Cambridge and Oxford NHS Blood and Transplant for the recruitment of study participants. We thank the Wellcome Sanger Institute Flow Cytometry, Sequencing, IT and Data Access facilities for their essential contribution to data generation and processing. We would like to acknowledge the enthusiasm and effort of Kiran Kumar Thurimella and Aman S. Patel in this project. We thank Emma Davenport for her feedback on the manuscript.

## Authors contribution

G.T. conceived the work; L.B.C., D.A.G., N.K. and G.T. designed the experiments; D.A.G., N.K., G.G., C.C. and D.J.S. performed FACS sorting, cell culture, DNA and RNA isolation and generated RNA-seq, ATAC-seq and ChIPmentation sequencing libraries; L.B.C. processed genotyping, ATAC-seq and ChIPmentation sequencing data; D.A.G. processed RNA-seq sequencing data; L.B.C. and D.A.G. analyzed and integrated the data; M.S. and I.D. performed drug tractability analyses; A.L. and D.R. recruited the blood donors; A.M. contributed to immune disease risk polymorphism enrichment analysis; K.A. contributed to the establishment of QTL analysis pipelines; B.S. and E.C.G. helped interpret the results; K.K. contributed to genotype imputation; N.S. contributed to the comparative analysis with naïve T cells; L.B.C., D.A.G. and G.T. wrote the manuscript.

## Materials and Methods

### Sample collection and Treg isolation

Lymphocyte cones were obtained with informed consent from donors at the NHS Blood and Transplant, Cambridge (REC 15/NW/0282) and from the NHS Blood and Transplant, Oxford (REC 15/NS/0060).

Leukodepletion cones were obtained from healthy adults of Caucasian origin. PBMCs were isolated using Lympholyte-H (Cedarlane Labs, <u>Burlington, Canada</u>) density gradient centrifugation. CD4+ T cells fraction of the PBMCs was obtained by negative selection using EasySep® Human CD4+ T Cell Enrichment Kit (Cat. no. 19052, StemCell Technologies, Vancouver, Canada), following the manufacturer’s instructions. Next, the CD4+ T cells were resuspended in the FACS staining buffer (2 mM EDTA and 0.5% FCS in PBS) at 10^8^ cells per ml. The cells were stained with the following antibody cocktail: anti-CD4-APC (30 µl/ml final volume, clone OKT4, Cat. no. 317416, BioLegend, San Diego, U.S.), anti-CD127-FITC and (30 µl/ml, clone eBioRDR5, Cat. no.11-1278-42, Thermo Fisher Scientific, Waltham, U. S.) and anti-CD25-PE (80 µl/ml, clone M-A251, Cat. no. 356104, BioLegend) for at least 30 min at RT in the darkness. The cells were washed copiously with FACS buffer and resuspended at 10^8^ cells per ml in full medium (IMDM, 10% FCS) and kept O/N at 4ºC. Immediately before sorting, the cells were stained with DAPI, to discriminate live and dead cells. The CD4^+^, CD25^high^, CD127^neg^ population corresponding to Treg lymphocytes was sorted out for the downstream assays (**Supplementary Figure 1**).

### Sample summary

For all donors we were able to extract their sex based on their genotype and for 113 of the 123 donors we had access to their age (**Supplementary Figure 2E**). The majority of the donors (78%) were genetically assigned males and were aged over 57 years of age.

### FACS staining

To verify the FOXP3 expression in the sorted Treg populations after sorting, the cells were stained for expression of CD4, CD25 and CD127 surface markers, and then stained with anti-FOXP3-BV421 antibody (5 µl/10^6^ cells, clone 206D, BioLegend) using the eBioscience™ Foxp3 / Transcription Factor Staining Buffer Set (Thermo Fisher Scientific), according to the manufacturer’s instructions. We observed that the sorted cells were on average 80% FOXP3 positive (**Supplementary Figure 2F**).

To define the proportions of memory and naive cells in the CD4^+^ population, an aliquot of 10^6^ cells after the CD4-enrichment were resuspended in 100 µl FACS buffer and stained with a cocktail of anti-CD4-APC and anti-CD127-FITC antibodies (3 µl each), anti-CD25-PE (8 µl) and anti-CD45RA-BV785 (4 µl, clone HI100, Cat. no. 304140, BioLegend), incubated at RT in the dark for at least 30 min., washed copiously with FACS buffer and analysed on BD Fortessa. The majority of the isolated Tregs were memory Tregs (median = 79%) (**Supplementary Figure 2G**).

### Culture and stimulation of isolated Tregs

Whole blood samples were obtained from ten healthy adults, aged from 22 to 39 years. Live regulatory T cells (CD4+ CD25high CD127low) were isolated as described in S*ample collection and Treg isolation*. Cells were grown in Iscove’s Modified Dulbecco’s Media (IMDM) (Life Technologies, Paisley, UK), supplemented with 10% human serum (HS), 50 U/ml penicillin and streptomycin (Life Technologies) and 100 U/ml recombinant human IL-2 and incubated at 37°C in a humidified atmosphere of 5% CO_2_. Cells were activated using PMA (5-10 ng/μl) with ionomycin (200 ng/μl) (Sigma-Aldrich) overnight (18 hours).

### SNP genotyping and imputation

A total of 551,839 genetic markers were genotyped using the Infinium® CoreExome-24 v1.1 BeadChip by Illumina. After SNP QC (MAF > 10%, SNP call rate > 95%, Hardy-Weinberg equilibrium (HWE) p-value < 0.001) we retained 243,820 variants in our dataset. Samples with call rate < 95% were removed from the analysis. After quality control per individual, the total genotyping call rate reached > 99%. We performed imputation using BEAGLE 4.1 with a reference panel comprising the 1000 Genomes Phase 3 ^58^ and the UK10K ^59^ samples (modelscale parameter = 2). Following imputation we required allelic R-squared (AR2) >= 0.8, HWE p-value < 0.001, and MAF > 10% in both the analysed cohort and in the reference panel, resulting in 4,640,249 variants in our final dataset. Of those, 512,320 were insertion-deletions (INDELs) and 608 were multiallelic polymorphisms. All genetic variant coordinates were lifted over to GRCh38.

Our samples clustered with the European populations included in the 1000 Genomes project (**Supplementary Figure 2H**). We removed 1 sample due to high relatedness (identity by state, pi_hat > 0.2). We used VerifyBamID v1.0.0 ^60^ with the genotype information along with all the functional genomics sequencing assays (see below) to verify no sample swaps were present in the final dataset.

### RNA-seq

For RNA-seq experiments, 0.5 × 10^6^ sorted Treg cells were washed with ice-cold PBS and resuspended in TRIzol (Thermo Fisher Scientific). After standard phenol/chloroform isolation step, the total RNA contained in the upper, aqueous phase was further purified with RNeasy Mini Kit (QIAgen, Hilden, Germany), according to the manufacturer’s instructions. The RNA libraries were constructed using KAPA RNA HyperPrep Kit (Roche, Basel, Switzerland), following a standard automated protocol. The libraries were multiplexed and sequenced at 75 bp PE on an Illumina HiSeq V4 to yield on average 57 million reads per sample.

### ATAC-seq

ATAC-seq was performed according to protocol ^61^, with following modifications. After sorting, the T cells were washed with ice-cold PBS and resuspended in sucrose buffer (10 mM Tris pH 8, 3 mM CaCl2, 2 mM MgOAc, 1 mM DTT, 0.32 M sucrose, 0.5 mM EDTA, 0.25% TritonX-100), followed by 5 min incubation on ice to isolate the nuclei. Isolated nuclei were washed once with 1x TD buffer (Tagment DNA Buffer, Nextera DNA Library Prep Kit, Illumina, U.S) and resuspended in 50 µl 1x TD buffer containing 2.5 µl of Tn5 enzyme (TDE1, Nextera). The reaction was carried out at 37ºC, mixing and then stopped by addition of 250 µl of buffer PB (MinElute PCR Purification Kit, QIAgen, Hilden, Germany). The DNA was then purified on MinElute columns according to the manufacturer’s instructions and eluted in 10 µl sterile ddH_2_O. The libraries were amplified using the NPM mix (Nextera PCR Master Mix from Nextera DNA Library Prep Kit) and Index adapters i7 and i5 (Nextera Index Kit, Illumina, U.S), according to the manufacturer’s instructions. The number of amplification PCR cycles for each sample was determined individually by performing a qPRC reaction of 7.5 µl aliquote of the mix with an addition of the EvaGreen dye (Biotium, Fremont, U.S.). The amplified libraries were SPRI purified (upper cut 0.5x, lower cut 1.8 x) on a Zephyr G3 SPE Workstation (PerkinElmer, Waltham, U.S.), multiplexed and sequenced at 75 bp PE on an Illumina HiSeq V4 to yield on average 112 million reads per sample.

### H3K4me3 and H3K27ac ChIPmentation-seq

The ChIPmentation-seq (ChM-seq) protocol was performed on 100,000 sonicated cells according to the protocol presented in Schmidl et al. 2015 and adapted to work with the iDeal ChIP-seq Kit for Histones (Diagenode, Liege, Belgium).

After sorting, the cells were resuspended in pre-warmed full medium (IMDM, 10% FCS) at 1-2 million cells per ml and allowed to recover in the incubator (37ºC, 5% CO_2_) for at least 30 min. The cells were then fixed by addition of formaldehyde to medium to a final concentration of 1% and 5 min incubation at 37ºC, followed by quenching with glycine for 5 min at a final concentration of 125 mM min at RT with mixing. The cross-linked cells were subsequently washed twice with ice-cold PBS and snap-frozen by immersion in liquid nitrogen.

0.5 × 10^6^ frozen cells were resuspended in 250 µl buffer iL1 with proteinase inhibitors cocktail (iDeal ChIP-seq Kit for Histones, Diagenode) and incubated for 10 min at 4ºC on the Bohemian wheel. The samples were then spun down, and resuspended first in buffer iL2 with proteinase inhibitors, then in iS1 with proteinase inhibitors, in both cases also for 10 min. at 4ºC. The cells were then sonicated in buffer iS1 using the Bioruptor® Pico sonication device (Diagenode) to achieve fragment sizes distribution below 3 kb.

Sonicated chromatin from 100,000 cells was used for an overnight immunoprecipitation reaction with 1 µg of antibody, either against H3K4me3 (Catalog No: 39915, Active Motif, Carlsbad, U.S.) or H3K27ac (Cat. no. C15410196, Diagenode).

The samples in deep-well plates were then washed twice for two minutes with 150 µl of each of the buffers: iW1, iW2, iW3 (iDeal ChIP-seq Kit for Histones, Diagenode) and then with 10 mM Tris pH 8. All the washes in this protocol were performed using an Agilent Bravo Automated Liquid Handling Platform (Agilent, Santa Clara, U.S.). After the second Tris wash, a ChIPmentation reaction on the beads was conducted following the protocol outlined in Schmidl. et al. Briefly, a mix containing 1 µl Tn5 from the Nextera kit was added to the beads and incubated for 10 minutes with vigorous mixing at 37ºC. Next, the reaction mix was removed using Bravo, and additional washes were performed, two with buffer iW3, followed by two washes with buffer iW4. The enriched DNA was eluted from the beads by incubation with 67 µl buffer iE1 (1h, RT, vigorous shaking). 3 µl of iE2 buffer were then added to each sample and the cross-linking was reversed by an overnight incubation at 65ºC in a thermocycler.

The DNA was then purified twice using SPRI beads at 1.6x ratio using a Zephyr G3 SPE Workstation. The libraries were amplified following the ATAC-seq library amplification protocol, but using NEBNext® High-Fidelity 2X PCR Master Mix (New England Biolabs, Ipswich, U.S.). Finally, the ChIPmentation libraries were sequenced to a depth of at least 13 million reads per sample and an average of 75 million reads per sample.

### RNA-seq data processing

Reads were aligned to the GRCh38 human reference genome using STAR ^62^ and the Ensembl reference transcriptome (version 87). Gene counts was performed using featureCounts tools from the subread package v1.5.1 ^63^ and only assigned reads were used for further processing (59.26% of reads were assigned). We excluded short RNAs and pseudogenes from the analysis. We quantile normalised the gene expression values and corrected for GC-content using the CQN method ^64^. We kept 12,059 genes with average count per gene across all donors greater than 25.

### Chromatin marks data processing

Reads were trimmed using skewer. Trimmed reads were aligned to the GrCh38 assembly of the human genome using bwa ^65^ and employing the mem algorithm. Multi-mapping reads and duplicated reads were removed using samtools ^66^. For ATAC-seq data, reads aligning to the mitochondrial chromosome were also removed. Only reads mapping to autosomes were maintained. A median of 30 million reads, 27 million reads and 40 million reads in the ATAC-seq, H3K4me3 ChM-seq and H3K27ac ChM-seq assays respectively, passed this QC. Peak calling was performed using MACS2 ^67^ independently on each donor for quality control purposes. For ATAC-seq peaks were called using the standard MACS2 model and specifying --nomodel --shift −25 --extsize 50 on fragment BED files (this is, both reads of a pair were merged into a single fragment). Prior to calling peaks from histone ChM-seq data, we merged a combined input reaching more than 223 million reads. H3K4me3 peaks were called using the standard narrow peak MACS2 model and an adjusted p-value of 0.01, specifying -f BAMPE --down-sample -q 0.01. H3K27ac broad peaks were called using the standard broad peaks macs2 model and an adjusted p-value of 0.01, specifying -f BAMPE --broad --down-sample --broad-cutoff 0.1 -q 0.01.

Samples with less than 10,000 peaks (median: ATAC 36,331, H3K4me3 22,815, H3K27ac 68,626), fraction of reads in peaks (FRiP) lower than 10% (median: ATAC 23%, H3K4me3 52.64%, H3K27ac 63.9%) (**Supplementary Figure 2C**), or, for ATAC-seq, an abnormal insert profile (defined as a ratio of short inserts (<150bp) over long inserts (>150) smaller than 1.5; average 2.03) were discarded. Additionally, the samples that did not cluster with the corresponding group in principal component analysis (considering log2 transformed number of reads in genomic bins of 10,000 bp, after normalization by library length) were discarded from further analysis. Finally, a total of 73, 88 and 91 individuals passed these filters for ATAC, H3K4me3 and H3K27ac samples, respectively. Sixty-two donors passed QC steps for all the tested omic layers (RNA, ATAC, H3K4me3 and H3K27ac).

In order to define a consensus set of peaks per chromatin assay, we performed a merged peak calling combining reads from all the donors. We downsampled each donor sample using samtools to 2 million fragments per ATAC-seq assay, 1.87 million read pairs per H3K4me3 and 1.86 million read pairs per H3K27ac assay in order to reach similar read counts to the sequenced inputs. We used the MACS2 parameters described above and specifying --keep-dup all. Then, to ensure a sufficient number of reads per peak, only ATAC-seq peaks with at least 10 reads in 80% of the samples, and ChM-seq peaks with fold enrichment >= 2 and adjusted p-value < 0.001, were maintained in the final set. The consensus sets were 39,642 ATAC-seq narrow peaks, 40,285 H3K4me3 ChM-seq narrow peaks and 34,457 H3K27ac broad peaks. The median length was 523bp, 794bp and 4501.5bp for the ATAC-seq peaks, H3K4me3 peaks and H3K27ac peaks, respectively. The median number of read pairs in each peak (calculated using featureCounts -p -C -D 5000 -d 50) per sample amounted 37.95 in ATAC-seq, 16.34 in H3K4me3 ChM-seq and 123.38 in H3K27ac ChM-seq. The peak overlap between the assays as calculated using bedtools intersect, the distance to the closest transcription start site (TSS) is shown in **Supplementary Figure 2B)**.

Genome browser data was constructed using the MACS2 -B flag and reads were normalised to signal per million. The fold-enrichment was calculated using the input background and finally bigwigs were constructed using bedGraphToBigWig command from the UCSC suite of tools ^68^. Coverage plots were generated using an adapted version of the wiggleplotr R Bioconductor package ^69^.

### Quantitative trait locus mapping (QTLs)

Prior to the QTL analysis we removed genes and peaks mapping to the MHC region (chr6: 20,000,000-40,000,000) and only kept the autosomal chromosomes. We used linear regression implemented in the QTLtools ^29^ software to map cis QTLs. For the gene expression we used a 500kbp cis-window around the gene, while for the three gene regulation assays we used a 50kbp cis-window around the defined peak. We used the “--permute 10000” to obtain permutation p-values for each gene and peak. As covariates we used the top 13, 30, 22 and 33 principal components that each explained up to 1% of the observed variance in the RNA, ATAC, H3K27ac and H3K4me3 layers, respectively. We picked the top most significantly associated variant for each gene or peak and used false discovery rate (FDR) correction to identify genes or peaks with at least one significant QTL at 5% FDR level.

To perform comparative analysis between Treg and naive T cell eQTLs and actQTLs we downloaded the naive CD4+ T cell RNA-seq and ChIP-seq datasets generated by the BLUEPRINT consortium ^24^ from the EGA archive (EGAD00001002671, EGAD00001002673) and processed the data using the same workflow as described above. We included the top 14 and 17 PCs for this dataset in the eQTL and actQTL analyses, respectively. We chose this dataset because these two cell types are closely related, the naive T cell dataset is of similar size (169 individuals) to the Treg dataset and all the individuals were of British origin. We used the following three criteria to define an eQTL and actQTL as cell type specific when comparing naive and regulatory T cells: (i) the gene was expressed in one cell type only or the peak was only present in one cell type (peak overlap was assessed using bedtools ^70^), (ii) the gene or peak was a significant QTL in one cell type (FDR ≤ 0.05) and not in the other (FDR > 0.2) and (iii) if the same gene or peak was a QTL in both cell types and the LD between the top QTL variant in regulatory T cells and any of the significant associated signals in naive T cells was lower than R^2^<0.2.

### Colocalization of QTL signals with immune disease GWAS

We used *coloc* v2.3–1 ^31^ to test for colocalization between molecular QTLs and GWAS SNPs listed in **Supplementary Table 1** and **2**. We included in our analysis the summary stats for the genome-wide association studies of 14 immune related diseases: allergic diseases (ALL; ^39^, ankylosing spondylitis (AS; ^71^, asthma (AST; ^72^, celiac disease (CEL; ^73^, multiple sclerosis (MS; ^74^, primary biliary cirrhosis (PBC; ^75^, psoriasis (PS; ^76^, rheumatoid arthritis (RA; ^77^, systemic lupus erythematosus (SLE; ^78^, type 1 diabetes (T1D; ^79^, vitiligo (VIT; ^80^, inflammatory bowel disease (IBD), Crohn’s disease (CD) and ulcerative colitis (UC) ^81^. We selected these diseases because they had more than 40 GWAS associated independent loci at p-value < 10^−5^. We also tested two non-immunological traits as controls, with a similar number of loci, type-2 diabetes (T2D; ^82^ and depression (DEP; ^83^. CD, UC and IBD were counted as a single disease when counting for number of colocalizing diseases per gene or peak. *Coloc* tests five hypothesis for colocalization. Hypothesis zero (H0) tests whether there is any association at all, PP.H1 and PP.H2 test whether there is an association with just one or the other study, PP.H3 tests whether the signal from GWAS and QTL is due to two independent SNPs, and PP.H4, test if the association between GWAS and QTL is due to a shared causal variant.

Prior to colocalization, we repeated the QTL mapping in chromatin features using a 500kbp window, to run *coloc* at a larger window. We ran *coloc* on a 400-kb region centered on each lead eQTL and chromQTL variant that was less than 100kb away from a GWAS variant (nominal p-value < 10^−5^). We only kept the colocalizations between QTLs and non-HLA GWAS loci if there were more than 50 SNPs tested. To claim a true colocalizing signal we required PP.H4 to be equal or greater than 0.8, the sum of the probabilities of having a significant signal in both studies to be equal or greater than 0.8 (PP.H3+PP.H4 ≥ 0.8) and the probability of the signal being due to a shared variant (PP.H4) was no less than 0.8 of the probability of having a significant signal in both studies (PP.H4/(PP.H3+PP.H4) ≥ 0.8) ^3^. In order to decrease the number of false positive findings in our Treg dataset, we focused on the colocalization results with common immune disease variants (MAF > 10%). All colocalizations between Treg QTLs and disease GWAS signals were manually verified and false colocalizations were discarded. For GWAS loci with 10^−5^ > p-value > 5 × 10^−8^, and colocalizing with Treg QTLs were verified in the original publications that there was a replication cohort and the final GWAS p-value was below genome wide significance.

### Gene expression variance deconvolution

To estimate the contribution of the genetic and epigenetic layers to the transcriptome variance we fitted a multivariate linear model to the expression of each gene. Therefore, the dependent variable in the model was the gene expression, while the independent variables were the genetic variants and/or chromatin features. We regressed out the PCs included in the analysis of each layer previous to model calculation. As described in de Bakker et al. ^84^, we performed an initial variable selection step by identifying the genetic variants or chromatin features in a +-150kb window that significantly correlated with gene expression (Spearman p-value < 0.05). In order to keep only independent variables in the set of predictors, in the instances where pairs of genetic variants or chromatin features were correlated (Spearman correlation > 0.4), we removed the variable with a lower correlation with gene expression. To determine the total variance explained we used the adjusted R^2^ of the model where we included all the independent variables from genetic variants and chromatin features The contribution of each individual layer, the genetic variants or chromatin features, or combination of the layers, was obtained by subtracting the R^2^ estimates of the models that excluded individual layers, or their combinations, from the R^2^ of the model with the total variance explained by all the layers.

### SNP functional classification

We retrieved evidence of functional relevance per genetic polymorphism in our colocalization results by integrating the four QTL analyses with the location of the histone modification or accessible chromatin enriched regions (peaks). We classified the genetic polymorphisms as “in peak” if they mapped in at least one of the peaks in the ATAC-seq, H3K27ac ChM-seq or H3K4me3 ChM-seq sets. Then, all the lead polymorphisms or their proxies (all polymorphisms showing R^2^ > 0.8 with the lead QTL signal) of significant eQTLs were classified as “eQTL”. Finally, all the lead polymorphisms of significant caQTLs (QTL in the chromatin activity layer, ATAC-seq), actQTLs (QTL in the histone acetylation modification layer, H3K27ac ChM-seq), and promQTLs (QTL in the histone promoter modification layer, H3K4me3 ChM-seq), or their proxies, were included in the chromatin QTL category. Figure 2D shows the number of polymorphisms included in the different categories, categories are mutually exclusive and each polymorphism is assigned only to the most functional one possible.

### LD loci definition and classification

A LD locus comprises a ±150kb window around the region defined by the lead and proxy variants (R^2^ > 0.8) for each GWAS signal that colocalizes with a Treg QTL. Loci with significant colocalization signals were classified as follows: i) Tier 1, at least one colocalization with an eQTL and at least one colocalization with a chromQTL; ii) Tier 2, at least one colocalization with an eQTL, no colocalization with chromQTL; and iii) Tier 3, at least one colocalization with a chromQTL (Figure 3 and **Supplementary Table 1** and **2**). We prioritize as functional variants in Tier 1 and Tier 3 loci which overlap with the regulated chromatin QTL peak.

### Known drug tractability evidence analysis

We used the Open Targets platform to extract information on drugs that support target-disease associations provided by ChEMBL ^85^. We retrieved the ‘known_drug’ evidence for all genes via the Open Targets API using the python client (Open Targets data release December 2018). For each target-disease-drug combination we determined the maximum clinical trial phase and the number of clinical trials in the Open Targets platform. In addition, information on target tractability available in the Open Targets platform (compiled by ChEMBL) was also retrieved for all genes.

### Differential gene expression analysis

RNA-seq reads were obtained and processed as described in *RNA-seq* and in *RNA-seq data processing*. Genes with at least 25 copies in at least three samples were kept, for a final table of 16,645 genes. Differential expression analysis was performed using DESeq2_.1.14 ^86^ Wald test, setting alpha at 0.05 and lfc at 1. Differentially expressed genes that mapped within a +/-150kb window from a chromQTL in Tier 3 loci were annotated as candidate stimulation specific eQTL genes.

### Data availability

Upon publication all data will become publicly available.

#### Software availability

Custom scripts and pipelines used for analysis in this study are accessible in the “Treg_Multiomic” repository: https://github.com/trynkaLab/

